# Identity Domains Reveal How Life Experiences and the Microbiome Shape Individuality and Uncover the Presence of Social Memory in Drosophila

**DOI:** 10.1101/2025.11.19.689229

**Authors:** Hadar Pozeilov, Elia Dayan, Rotem Yehuda, Mali Levi, Galit Shohat-Ophir, Oren Forkosh

## Abstract

Personality is comprised of enduring traits that shape behavior across contexts and over time. Yet outside humans, research often equates individuality with personality, focusing on how one animal differs from another in each behavior. This overlooks the latent space that spans an animal’s behavioral repertoire, how it is shaped by innate tendencies and prior experiences, and how it differs from transient factors. Here, we bridge this gap by integrating a data-driven framework with unbiased trait interpretation via large language models, revealing how social, sexual, and physiological conditions systematically alter the expression of personality-like traits in *Drosophila melanogaster*. This approach yielded four identities: space-use (exploration), time-use (social investment), avoidance, and aggression. Large language models produced strong consensus on the interpretation of some of these axes and divergence on others. These dimensions proved stable within individuals yet shifted predictably following mating, isolation, and microbiome manipulation, often in sex dependent ways. Notably, flies behaved differently toward familiar and unfamiliar peers, revealing a capacity for social memory. These findings bridge temperament and personality, showing how life history reshapes baseline tendencies into individualized profiles within constrained dimensions. They establish *Drosophila* as a tractable system for uncovering the biological logic of personality and advancing cross-species principles for individuality research.

**Significance Statement:** Personality is often treated as a hallmark of complex brains, yet we show that fruit flies express stable behavioral identities with systematic, experience driven plasticity. Using a mathematical framework, we resolve four stable traits and map how sex, social experience, and the gut microbiome shift their expression. We further demonstrate social recognition in flies: they treat familiar cage mates differently from newcomers over repeated encounters, evidencing social memory, a striking result given their simple nervous system. Finally, we introduce large language models to provide objective, reproducible interpretations of trait dimensions, reducing human bias. These advances establish a general, quantitative approach to personality and social behavior in a tractable genetic model.

## Introduction

Evolution shapes living systems through two seemingly contradictory forces: the stringent optimization of traits through natural selection and the persistent maintenance of variation within populations. As selection drives organisms toward adaptive peaks in their fitness landscapes, it simultaneously preserves a diverse spectrum of individual variants. This interplay between optimization and variation extends beyond genetic inheritance through phenotypic plasticity (1); throughout our lives, behavior is shaped by experience, with neuroplasticity shifting the balance of action selection and generating an additional layer of adaptive diversity that operates within, and sometimes transcends, inherited predispositions (2).

The intricate relationship between temperament and personality, as defined in human psychology, illustrates this evolutionary dynamic (3). Temperaments are biologically rooted behavioral predispositions present from early development, providing the foundation upon which personality emerges through continuous interaction with environmental experiences. Crucially, both temperament and personality vary along specific, constrained dimensions rather than showing random variation across all possible behavioral traits (4). In humans, for example, the core dimensions of personality are captured by the Big Five traits, commonly abbreviated as OCEAN: openness to experience, conscientiousness, extraversion, agreeableness, and neuroticism. The existence of these consistent dimensions suggests that natural selection has shaped which aspects of behavior can productively vary between individuals, permitting beneficial diversity while maintaining core adaptive functions (5).

While the evolutionary utility of personality traits is well-established, a fundamental question remains: how do individuals balance adaptability with consistency across life? One way to frame this question is by analogy to the evolutionary strategy of bet hedging: lineages facing unpredictable environments spread risk by generating variable phenotypes across time or offspring, thereby enhancing long-term survival (6, 7). By extension, stable individual differences may reflect a similar principle, with variability serving as a hedge against environmental uncertainty (8–14). Freund’s work illustrates this idea, showing that genetically identical mice housed together in enriched environments developed distinct behavioral patterns over time, with divergent exploration correlating with hippocampal neurogenesis (9). This provides evidence that developmental processes beyond genetic programming contribute to stable behavioral variation.

As personality research has expanded beyond humans to a broad range of animal species (15–20), it has become clear that stable individual differences are a common feature of behavior across the animal kingdom (21–23). Whereas human personality is commonly assessed via self-report, animal personality is inferred from consistent patterns of behavior across time and context (15). Field and laboratory studies reveal persistent differences in boldness, aggression, and sociability, among others (16, 19, 24).

Defining personality, however, remains challenging. Individuality can be measured directly from behavior, but personality is more abstract, aiming to capture the underlying trait dimensions that organize this variation (23). Although researchers generally agree that consistent responses to similar cues are adaptive, they differ in how they conceptualize and label these traits. Terms such as *temperament*, *behavioral syndromes*, *coping styles*, and *predispositions* are often used interchangeably, reflecting a lack of consensus about what constitutes personality in animals (25–30).

Beyond terminology, methodological issues complicate the field. Many studies rely on narrow or subjective behavioral readouts and are interpreted through an anthropocentric lens. For example, behavior in the light–dark box for mice is commonly taken as a proxy for anxiety, yet the same actions could reflect curiosity or exploration. Such ambiguities can mask true dispositions, as when some mice classified as “bold” in the elevated plus maze were later found to be blind (31). These challenges highlight a central problem: behavior is the expression of underlying traits, not a direct proxy for them. The aim of personality research is therefore not merely to catalog differences among individuals but to identify the trait dimensions that generate them.

This motivates the need for systematic and unbiased frameworks that move beyond assay-specific interpretations. Personality can be formalized as positions along behavioral dimensions rather than as isolated measures, a perspective recently realized through a data-driven approach known as identity domains (23). Using Linear Discriminant Analysis (LDA), this framework captures trait dimensions that maximize variability between individuals relative to variability within each individual and that remain stable over time. By integrating information across many behaviors, identity domains capture the structure of individuality in a species-agnostic way. First applied to mice, the framework has since been extended to other species, underscoring its potential as a comparative scaffold for studying personality across taxa (30).

The lack of a unified framework is further amplified by a strong taxonomic bias, with research on personality variation concentrated in vertebrates while invertebrates remain comparatively neglected (32, 33). Despite their diversity, ecological importance, and experimental tractability, invertebrates are often dismissed as behaviorally simplistic (32), a view that obscures opportunities to probe the evolutionary origins and mechanisms of personality across the tree of life.

Eusocial insects, including ants, bees, wasps, and termites, are especially important in this regard (34–36). Although invertebrates, their colonies operate as integrated ״superorganisms״, displaying a level of organization and division of labor that parallels the complexity often associated with vertebrates (37–40). Whereas colony members were once considered interchangeable, recent work reveals consistent between-individual differences across eusocial species (34, 41–43). In honeybees, high-resolution behavioral profiling and gene-regulatory analyses uncovered a continuum of worker phenotypes with molecular correlates (44) and stable scouting tendencies (45). In ants, morphological subcastes align with spatial organization and task allocation (46, 47). In wasps, five distinct personality-like traits separate along social and non-social dimensions (48). Beyond genes and epigenetics, the gut microbiome modulates behavior in social insects: manipulating symbionts in bees alters foraging intensity, linking microbial composition to task allocation and personality-related behavior (49–53).

Living in a colony also requires the ability to recognize others. This capacity can support cooperation, cohesion, and defense against intruders, and in most social insects it is mediated by colony-specific chemical signatures that serve as identity cues (54, 55). These cues are typically blends of cuticular hydrocarbons shared among nestmates, forming a unified “Gestalt” colony odor (54). Their importance is evident in honey bees, where guard bees assess incoming individuals at the hive entrance, and in invasive Argentine ants, where reduced variation in these cues has led to the formation of vast supercolonies with little aggression between nests (55, 56). Recognition systems vary in specificity: while many species primarily distinguish colony members from outsiders, others extend this ability further. Paper wasps, for instance, can discriminate among individuals using unique facial patterns, providing striking evidence of individual recognition in an insect society (57, 58). Recent work has begun to uncover the mechanisms behind these systems, from specialized chemosensory organs in ants that detect foreign scents (59) to the genetic basis of olfaction and social behavior (60), and even colony-specific microbiomes that help define group identity in honey bees (61).

At the same time, eusocial colonies are organized so that behavior and fitness consequences are aligned with colony needs, where reproduction is centralized and competition among nestmates is minimized (35, 36). This makes eusocial systems invaluable but less representative of the selective pressures existing across many species in the animal kingdom, where individual reproduction and survival are the primary targets of selection.

As a non-eusocial species with a simpler, smaller brain, *Drosophila* reinstates individual-level competition for resources and mates, removing the confounds of caste and age polyethism. Recent work in *Drosophila melanogaster* shows that stable, between-individual differences also emerge in solitary contexts where no colony structure enforces role differentiation; strikingly, such differences appear even among genetically identical, co-reared flies (62, 63). High-throughput behavioral pipelines reveal consistent variation across multiple behaviors that organize into behavioral syndromes, correlated traits that persist within individuals across contexts (64–66). Targeted neural and transcriptomic manipulations can selectively shift specific axes of this variation, indicating a mechanistic basis for stable, inter-individual differences even in the absence of genetic or environmental heterogeneity (66–68). Together, these findings establish *Drosophila* as a powerful model for investigating how individuality arises, offering a bridge between insights from eusocial insects and research in vertebrates, including humans.

Moreover, behavior is embedded in a social context, where group dynamics and hierarchies shape trait expression in ways often missed by assays of isolated individuals (69–71). In many species, including *Drosophila*, social status influences access to mates, food, and other resources (72–76), driving divergent behavioral strategies across ranks. Such interactions not only affect immediate behavior but can also produce lasting shifts in personality traits as individuals adapt to social challenges (77).

In this study, we investigate the basis of individuality in the fruit fly *Drosophila melanogaster* and uncover a dual structure in behavior. Flies exhibit remarkably consistent baseline patterns that form a behavioral fingerprint similar to temperament, yet specific life events such as mating produce significant and predictable shifts. By applying Linear Discriminant Analysis to high-resolution behavioral data, we defined identity domains, dimensions of variability that remain stable within individuals while distinguishing them from one another. These domains reveal how sex, social interactions, and microbiome composition influence the balance between behavioral consistency and adaptability. Our findings link temperament-like baselines with personality-like plasticity, showing how life experiences reshape inherent tendencies to generate unique behavioral profiles. We further demonstrate that flies behave differently when housed with familiar compared to unfamiliar conspecifics, suggesting the presence of social memory. Together, these results establish *Drosophila* as a powerful model for dissecting the mechanisms of personality and provide insights into the biological foundations of personality-like traits, even in organisms with relatively simple nervous systems.

## Results

### Quantifying Individual Behavioral Variation and Stable Traits in Flies

To quantify stable behavioral traits in *Drosophila melanogaster*, we employed an integrated behavioral profiling system that combines automated tracking with computational phenotyping (Fig. 1A). Our experimental design systematically varied early life experiences across male and female flies, including different levels of social enrichment and sexual interaction - factors previously demonstrated to influence fly physiology and behavior (53). We conducted all behavioral measurements in group settings to capture social dynamics, as these environments elicit diverse behavioral patterns through continuous interaction between individuals.

**Figure 1.**
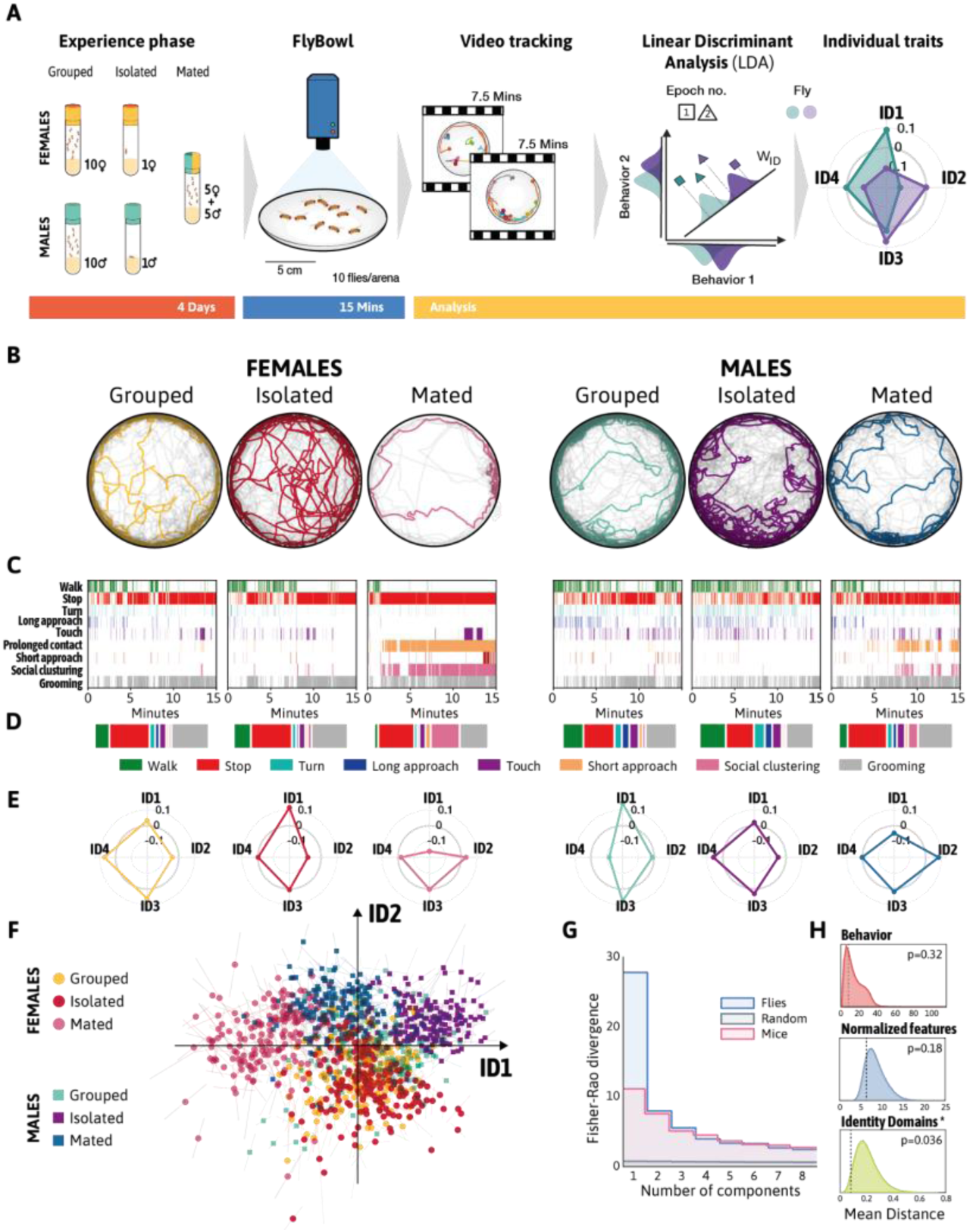
Emergence of Individual Differences in Fly Behavior. (A) Experimental design and analytical workflow. Newly flies were assigned to three social conditions: same-sex groups (10 flies), isolation (single flies), or mixed-sex groups (5 nd 5 females). After 4 days, flies were placed in same-sex groups of 10 in the FlyBowl apparatus for 15-minute behavioral ngs that were divided into two epochs. Video tracking generated 39 behavioral features, comprising the “Behavioral The Identity Domains method was applied to normalize behaviors by rank, transform them to normal distributions, and them via linear discriminant analysis onto a subspace optimized for individual differentiation while maintaining within-al consistency between epochs. This analysis yielded characteristic trait dimensions for each fly. (B-D) Representative ral signatures across early-life conditions. (B) Representative individual fly trajectories (colored) shown against group r trajectories (gray). (C) Corresponding ethograms displaying nine key behaviors: walking, stopping, turning, far ch, touching, prolonged contact, near approach, social clustering, and self-grooming. (D) Summary of behavioral on, showing the proportion of time spent in each behavior. (E) Individual trait profiles visualized using radar plots, ng the four most significant Identity Domains (ID1-ID4) for flies shown in B-D. (F) Two-dimensional ID space depicting sus ID2. Females (circles) and males (squares) are shown with condition-specific colors: grouped females (yellow), females (red), mated females (pink), grouped males (light green), isolated males (purple), and mated males (blue). The present the change in individual measurements across temporal epochs. (G) Traits ranked by their Fisher-Rao divergence representing the ratio of between-individual to within-individual variance and serving as a measure of trait reliability as ality indicators. The plot compares fly IDS (blue) with mouse IDs (pink), and a control shuffled dataset (gray). (H) Stability ison of raw behavioral features ranked features, and IDs across the two 7.5-minute epochs within the 15-minute test The stability was measured using the Euclidean distance between the two epochs. The p-values for each stability test played in each panel and were calculated by comparing the empirical results against shuffled data. Only the Identity s analysis yield a significant self-similarity (P=0.036).

For four days prior to the test, male and females were reared under the following conditions: one cohort of flies were raised in same-sex groups of 10 individuals, the second cohort raised from eclosion in complete social isolation (isolated), and the third cohort was raised in mixed-sex groups of 5 males and 5 females, allowing for voluntary and uncontrolled mating events (mated). After the experience phase, flies were introduced into circular arenas, the FlyBowl (54, 55), in same-sex groups of 10 and recorded for 15 minutes (Fig. 1A).

The position of each fly was tracked, generating data for 39 behavioral features, including movement velocities, relative positions, and 10 complex behaviors classified using a machine-learning-based system (JAABA; Supplementary Table 1) (55). The flies’ pre-test conditions resulted in distinct patterns of activity levels and action selection within the test arenas. Figures 1B-D illustrate these cohort-specific behavioral dynamics through representative locomotor tracks and ethograms, while Supplementary Figure 1 shows the temporal evolution of mean behavioral metrics for each experimental group.

The emergent behavioral responses during test reflect differences in the motivation state of group members, various degrees of activity levels, and sensory sensitivity to interacting flies, which could differ across the six cohorts. For example, the behavior of socially raised flies is expected to represent a snapshot of established relationships formed within the group during the experience phase early life experience. In contrast, the behavior of solitary flies is presumed to reflect the initial interactions of flies encountering others for the first time (54).

To isolate intrinsic, stable traits that distinguish one fly from another - independent of transient effects such as arena habituation - we turned to the Identity-Domain (ID) framework (23). In our implementation of the method, each experiment is first divided into two equal-duration epochs to minimize the influence of short-term factors. Within each epoch, raw values for every behavioral metric are converted to ranks, capturing each fly’s relative standing rather than its absolute level; thus, temporal changes in behavior do not mask consistent individual differences. These ranks are then Gaussian normalized to place all variables on a common scale. The normalized data from both epochs are entered into linear discriminant analysis (LDA), which derives linear combinations of behaviors that maximize between-individual variance relative to within-individual variance. These discriminant directions define the ID space - a concise set of trait axes on which each fly’s coordinates capture its stable, personality-like profile. Applied to our full dataset of 1,080 flies, this pipeline yielded a robust, interpretable map of individual behavioral identities (Fig. 1E–F).

The Fisher-Rao divergence (DFR), defined as the ratio of between-individual to within-individual variance, serves as a “personality-likeness” score that allows us to rank behavioral traits obtained through the ID method and determine their significance. This metric enables a quantitative comparison of behavioral subspaces, with higher DFR values indicating traits that better capture consistent individual differences. Figure 1G presents the ranking of the flies’ IDs by their DFR values, alongside data from mouse IDs and a control dataset with shuffled identities. The first fly ID exhibited substantially higher discriminative power than its mouse counterpart, while the second, third, and fourth IDs were comparable across species. Based on the previously suggested threshold of DFR > 4 (Forkosh et al., 2019), the first four fly IDs were classified as significant.

To test whether Identity Domains (IDs) reflect intrinsic behavioral traits, we assessed their stability over time by comparing each fly’s ID scores across the two experimental epochs (Fig. 1H, bottom). For comparison, we performed the same comparison using the original 39 behavioral features (Fig. 1H, top) and again after applying rank normalization to those features (Fig. 1H, middle). While both raw and rank-normalized features showed no significant consistency across epochs, ID scores remained significantly stable over time, indicating that they capture trait-like, rather than transient, aspects of behavior.

To investigate the biological relevance of the IDs, we first examined their statistical relationships with the 39 behavioral features extracted from the tracking data (Fig. 2). Each trait is visualized using a “tree graph” that displays the 15 features most strongly correlated with a given Identity Domain. In these graphs, bar length represents correlation strength and direction, while bar color indicates the behavioral category. This visualization allows a compact overview of which behaviors are most strongly associated with each ID (Fig. 2).

**Figure 2.**
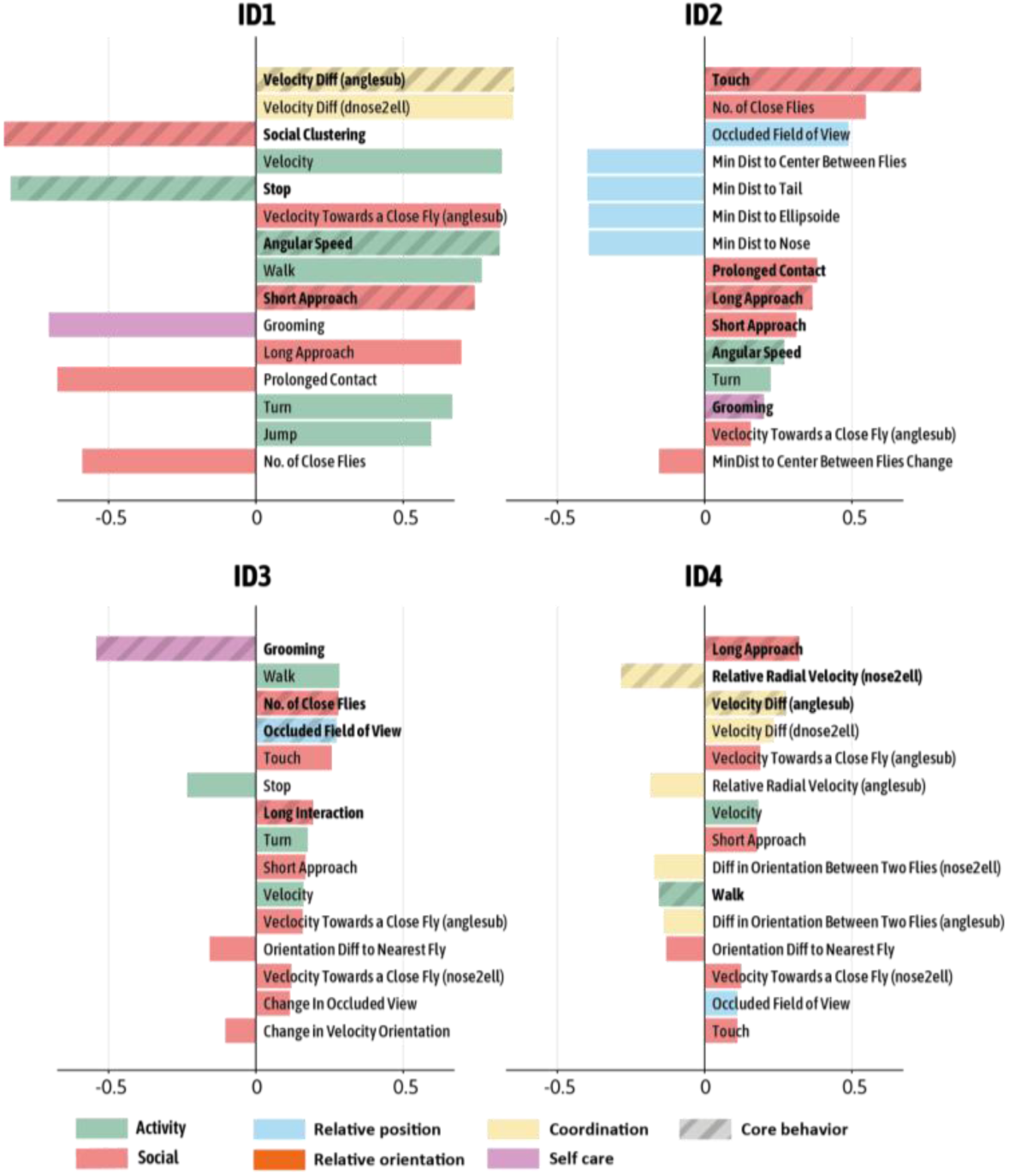
Mapping Behavioral Features to Identity Domains (IDs). This figure illustrates the Pearson correlations between 39 behavioral features and the four most significant IDs identified in the previous analysis. Behavioral features are grouped into six categories: activity, social, relative position (to conspecifics), relative orientation, coordination, and self-care. Each subplot displays a tree plot for one ID, showing the 15 most significant correlations. Bar length indicates correlation strength and direction (x-axis), while bar color denotes the behavioral category. Features are sorted from top to bottom by the absolute value of their correlation with the corresponding ID. To assess which features contribute uniquely to each ID, we also applied Lasso regression; features overlaid with a striped pattern indicate non-zero regression coefficients, highlighting their distinct contribution to trait identity.

To interpret and label the IDs derived from our behavioral data in an unbiased and reproducible manner, we employed multiple Large Language Models (LLMs) - including OpenAI’s ChatGPT 4.5, Google’s Gemini 2.5 Flash, and Anthropic’s Claude 3.7 Sonnet (see Table 1). By involving multiple models, we aimed to enhance the objectivity and robustness of trait labeling, reduce reliance on human interpretation, and mitigate model-specific biases. Each LLM independently analyzed the correlations between observed behaviors and the latent traits captured by the IDs, drawing on patterns from prior scientific literature to propose candidate labels. This process yielded strong consensus for ID1 (“Activity”) and ID2 (“Sociability”), whereas interpretations for ID3 and ID4 varied more substantially across models, reflecting greater ambiguity in their behavioral associations. The third trait, ID3, appeared to be related to social avoidance or generalized anxiety. The variability in labeling for ID3 and ID4 parallels their lower Fisher-Rao divergence values (Fig. 1G), suggesting these traits are inherently less discriminative across individuals. Nonetheless, as shown in subsequent analyses, even these less clearly defined dimensions proved informative for distinguishing experimental groups and condition-specific behavioral modulations.

**Table 1:** Interpretations of the Traits using Large Language Models. The table summarizes the traits the LLMs assigned to each ID based on a table of correlations between each trait and the meaningful behaviors. We used three LLMs: OpenAI’s ChatGPT 4.5, Google’s Gemini 2.5 Flash, and Anthropic’s Claude 3.7 Sonnet.

|  | ChatGPT 4.5 | Gemini 2.5 Flash | Claude 3.7 Sonnet |
| --- | --- | --- | --- |
| **ID1** | Activity/Aggression | Active/Exploratory | Activity/Exploratory |
| **ID2** | Sociability | Sociable/Interactive | Sociability/Social Interaction |
| **ID3** | Anxiety/Avoidance | Reclusive/Less Active | Social Tolerance/Gregariousness |
| **ID4** | Exploration/Curiosity | Unclear/Less Defined | Social Initiative/Assertiveness |

Building on the LLM-based trait labeling, we next examined the specific behavioral patterns that define each ID. We found that ID1 is characterized by high activity levels, increased velocity, and locomotion-related behaviors, negatively correlating with stationary behaviors such as stopping, grooming, and social clustering. Flies scoring high in this trait actively explore their environment, demonstrating significant mobility and frequent directional changes. The second dimension, ID2, represents time investment in social interactions. Flies scoring high in this dimension exhibit prolonged and substantial social engagement over time. ID3 is primarily characterized by high levels of grooming behavior and negatively correlates with active locomotion, proximity to other flies, and tactile interactions. Flies with high ID3 scores display increased grooming, reduced physical activity, and minimal social interactions. Specifically, they tend to avoid extensive walking, maintain greater interpersonal distances, and infrequently initiate physical contact. The fourth trait, ID4, is marked by high levels of long-distance approaches, increased velocity toward other flies, and shows lower differences in velocity directions relative to the nearest animal, behaviors consistent with chases and territorial aggression. Finally, while the observed traits share similarities with personality dimensions reported in other species, we emphasize that these findings capture behavioral individuality on relatively short timescales.

### Sex and Life History as Key Drivers of Behavioral Trait Variation

To examine the newly defined identity dimensions, we mapped the distribution of individual male and female flies raised in same-sex groups across the trait space. We used this basic life-history condition as a baseline to compare other experimental conditions and to identify intrinsic differences between sexes. Here, the position of individuals is visualized by projecting their relative scores onto a two-dimensional trait space (Fig.3A-C). Comparing the distribution of socially raised males and female flies within the space formed by combinations of trait dimensions revealed inherent sex-specific differences in this seemingly simple and uniform condition in 3 out of the 4 traits. Specifically, male flies exhibit overall higher scores in space-use (ID1) and time-use (ID2) as well as in aggression (ID4), suggesting that the expression of these traits is modulated by sex, whereas the avoidance trait (ID3) is similar between the sexes. The distribution of individuals from both sexes across these trait maps highlights the ability of the four identity dimensions to effectively distinguish between individuals.

**Figure 3.**
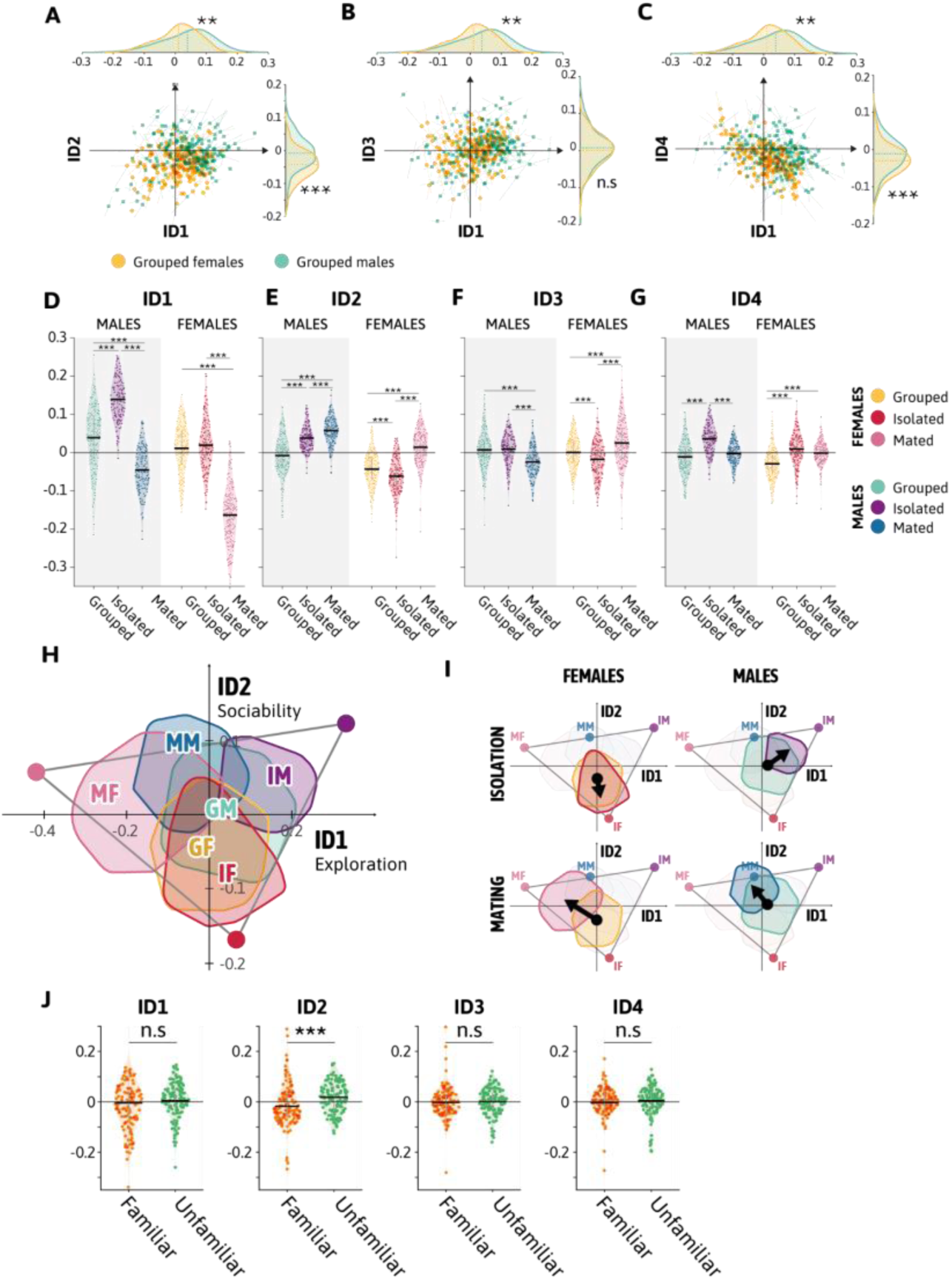
Early Life Experiences Shift the Expression of Individuality Traits. In all panels, statistical significance is indicated by *, **, and ***, for p < 0.05, p < 0.01, and p < 0.001, respectively; “n.s.” denotes non-significant differences (p > 0.05). A-C. Identity Domain (ID) spaces of the flies, showing pairwise combinations of ID1, ID2, ID3, and ID4 for grouped females (yellow circles, n = 180) and grouped males (green squares, n = 180). Within-fly variability across epochs is visualized as colored outlines around each marker (yellow for females, green for males). Kernel density estimates are shown along each ID axis, separately for each sex and color-coded accordingly. D-G. Trait differences across conditions. These panels show violin plots illustrating the distributions of ID values for grouped, isolated, and mated (Males: n = 180 each; Females: n = 180 each). Distributions are shown separately for each sex. One way permutation tests were performed to assess differences between conditions for each sex. H. Clustered ID space across all six conditions, based on ID1 and ID2. The trait space forms a triangular structure defined by three archetypes. Clustering was performed using a 2D kernel-based method. Each cluster is labeled using a two-letter acronym: the first letter indicates life experience (G = grouped, I = isolated, M = mated); the second indicates sex (F = female, M = male). For example, “IM” refers to isolated males. I. “Force maps” illustrating directional trait shifts. Arrows represent changes in cluster centers from the baseline condition (grouped males or females) to other life experiences, reflecting the directional influence of early-life conditions. J. Trait differences between flies housed in their original familiar groups (n = 130) and in mixed groups of unfamiliar individuals (n = 130). Violin plots display the distributions of Identity Domain scores, with a significant difference observed only in sociability (ID2).

To parse out the interplay between adaptive responses to various motivational states and the expression of individuality traits, we extended the analysis to include all six cohorts, accounting for the effects of social and mating experiences (Fig. 3D-G). We found that the degree of social enrichment and prior mating history shifted the positions of flies to distinct locations across the trait map. Analyzing individual fly scores for space-use (ID1) revealed that sex, along with the degree and type of prior social enrichment, had distinct effects on the relative positions of flies along this axis, with solitary males scoring the highest and mated females scoring the lowest (Fig. 3D). In addition to the distinct distribution of male and female flies along this dimension, we documented discrete distribution patterns associated with social and mating status within each sex. In male flies, social isolation heightened trait scores, group living reduced these scores, and mating further decreased scores related to spatial exploration. A similar trend exists in females, were mating dials-down the scores of ID1 (Fig. 3D). The differences in space-use scores are further evident when examining the density of representative trajectories associated with the different cohorts, highlighting the strong impact of prior experiences known to modulate motivational states on behavioral variation (supp Fig. 2). These findings highlight the ability of this trait to effectively distinguish between males and females under the same conditions, as well as to capture differences across varying social and mating experiences within each sex. Socially raised male and female flies differed significantly in their spatial exploration patterns (supp Fig. 2), reflecting inherent dimorphic differences in foraging-related behaviors. However, the overall trajectory of space-use was similar across sexes, transitioning consistently from higher scores in solitary individuals, to intermediate scores in socially raised flies, and to the lowest scores in mated flies. This indicates that while the expression of this trait is modulated by sex, the mechanisms governing responses to social and mating experiences are largely conserved.

Exploring the interplay between life history and time investment in social engagement (ID2) demonstrated the power of this trait in discriminating between individuals, revealing the extent by which different elements dial up and down the scope of sociability (Fig. 3E). Comparing the distributions of male and female that share similar life histories along this scale, reveals a shift towards higher scores in male flies, suggesting that this trait has a capacity to capture sex specific differences in the expression of social time-use (Fig. 3E). Examining each sex separately uncovered further complexity: solitary and mated males score higher than socially raised males, a finding also supported by analyzing spatial densities of flies across the arena (Supp Fig. 2). Although solitary and mated males exhibited elevated scores, their distribution patterns differ, suggesting that this trait captures different types of social engagements. In contrast, female cohorts exhibited different responses to the same conditions, with solitary females scoring lowest and mated females scoring highest, indicating that experience-dependent shifts differ markedly between sexes. Specifically, while social isolation enhances sociability in males, it significantly reduces sociability in females. This gender-specific response is especially notable, as detailed behavioral comparisons between solitary and socially raised females across multiple behaviors failed to detect significant differences (Supp. Fig. 1).

Analyzing individual positions along ID3 produced intriguing and somewhat unexpected findings. Mated females had the highest scores, whereas mated males had the lowest scores, highlighting a pronounced sexual dimorphism under the same condition (Fig. 3F). Among males, only mating influenced scores, while differences in social conditions did not significantly alter male positioning along this dimension. Conversely, female cohorts exhibited distinct distributions: mated females had the highest scores, solitary females had the lowest (resembling mated males), and socially raised females had intermediate scores (Fig. 3F). It is also notable that solitary males and females differed in their distributions along the avoidance dimension, emphasizing further sex-specific variability in response to social and mating experiences (Fig. 3F).

Lastly, we analyzed the patterns associated with the fourth identity dimension ID4. This dimension likely captures agonistic behaviors related to aggression, defense, or competition over resources. Consistent with previous studies linking social isolation to increased aggression (56–60), isolated males scored highest among male cohorts, indicating that a lack of social interaction enhances competitive behaviors, supporting the hypothesis that ID4 captures individual variability in agonistic interactions. Among males, solitary individuals showed higher scores than both mated and socially raised males, with socially raised males having the lowest scores (Fig 3G). Females displayed a similar pattern: solitary females scored higher than socially raised females, who again exhibited the lowest scores within female cohorts. Notably, although the overall aggression scores were lower in females, the relative distribution patterns across solitary, socially raised, and mated groups were parallel between sexes, emphasizing a shared underlying response mechanism where social experience consistently reduces aggression scores (Fig 3G).

Examining the ranking of female flies across the four new IDs reveals an intriguing pattern: each cohort is most distinctly separated by a specific ID (Fig. 3D-G). Mated females show the greatest differences in ID1 and ID3, solitary females rank lowest on the social time-use trait (ID2), and grouped females are uniquely characterized by the lowest scores on the ID4 scale. Together, these findings illustrate how social and sexual conditions dynamically modulate individuality and behavioral traits, with sex-specific effects and the interplay of life experiences acting as key drivers of variation.

Next, focusing on the two most prominent traits, we identified three distinct archetypes that define the boundaries of behavioral variation (Fig. 3H): isolated males (IM), isolated females (IF), and mated females (MF). These archetypes were derived using Pareto optimality analysis, which revealed that the trait space is bounded by a triangle with the archetypes as its vertices (61), aligning with findings in other species (23). In this context, archetypes represent individuals with extreme trait combinations that define the edges of the behavioral space.

Analysis of directional “forces” within the trait space defined by combinations of ID pairs revealed intersectional patterns shaped by sex and motivational states (Fig. 3I). In males, isolation and mating drove the cluster (GM) in opposite directions along the ID1 axis (space-use), but in the same direction along ID2 (time-use). In females, mating elicited similar effects along the ID2 axis (time-use), whereas isolation did not induce a shift, highlighting a sexually dimorphic response to social deprivation, and the role of sex-specific social dynamics in shaping individual behavioral trajectories.

Our findings demonstrate that the trait space defined by the four IDs effectively differentiates individual flies, assigning each a unique combined score. Furthermore, social and sexual conditions influence the positioning of individuals within the trait space while preserving differences among individuals within the same cohort.

Given the significant differences between solitary and socially raised flies, we next examined how social context influences the expression of identity traits by analyzing the spatial distribution of socially raised flies tested with either familiar or unfamiliar flies (Fig. 3J). Male flies were raised in groups of 10 individuals for four days and subsequently tested in two contexts: groups comprising the original, familiar individuals or mixed groups consisting of unfamiliar flies from different vials. Despite these different social contexts, the trait distributions remained consistent for most identity dimensions except sociability (ID2). This consistency was further supported by the absence of significant differences in overall behavioral features (Supp. Fig. 1C), indicating the robustness and stability of these personality-like traits despite changes in social conditions. Intriguingly, the distribution along the sociability dimension revealed that flies interacting with unfamiliar individuals exhibited higher scores, suggesting that flies possess social recognition capabilities and invest more time in social interactions when encountering new flies. This underscores the unique sensitivity and analytical power of the social time investment dimension.

### Microbiome and Social Stress Shape Personality Trait Dynamics in Drosophila

Having established a framework to explore stable behavioral traits among individuals and their dynamic responses to motivational states, we next broadened our analysis to investigate how negative experiences influence the expression of identity traits. Specifically, we examined whether being raised with aggressive flies - an experience analogous to social defeat in rodents (62) - impacts the expression of personality-like traits in non-aggressive flies. Wild-type (wt) male flies were cohabitated with hyper-aggressive males (generated by knocking down Cyp6a20; (63) for three days (Fig. 4A). At the end of this experience, we assessed the behavior of the wt flies in groups of ten. In contrast to socially defeated rodents, which typically exhibit low level of sociability, flies raised with hyper-aggressive individuals demonstrated significantly increased scores along the social axis (ID2); reduced activity scores (ID1; Fig. 4B); a slight but significant elevation along the anxiety/avoidance dimension (ID3), which it is in agreement with socially defeated rodent; and a marked reduction in ID4. The unique combination of their four IDs suggests that being raised in an aggressive environment may facilitate higher sociability scores, possibly as a strategy to mitigate social tension.

**Figure 4.**
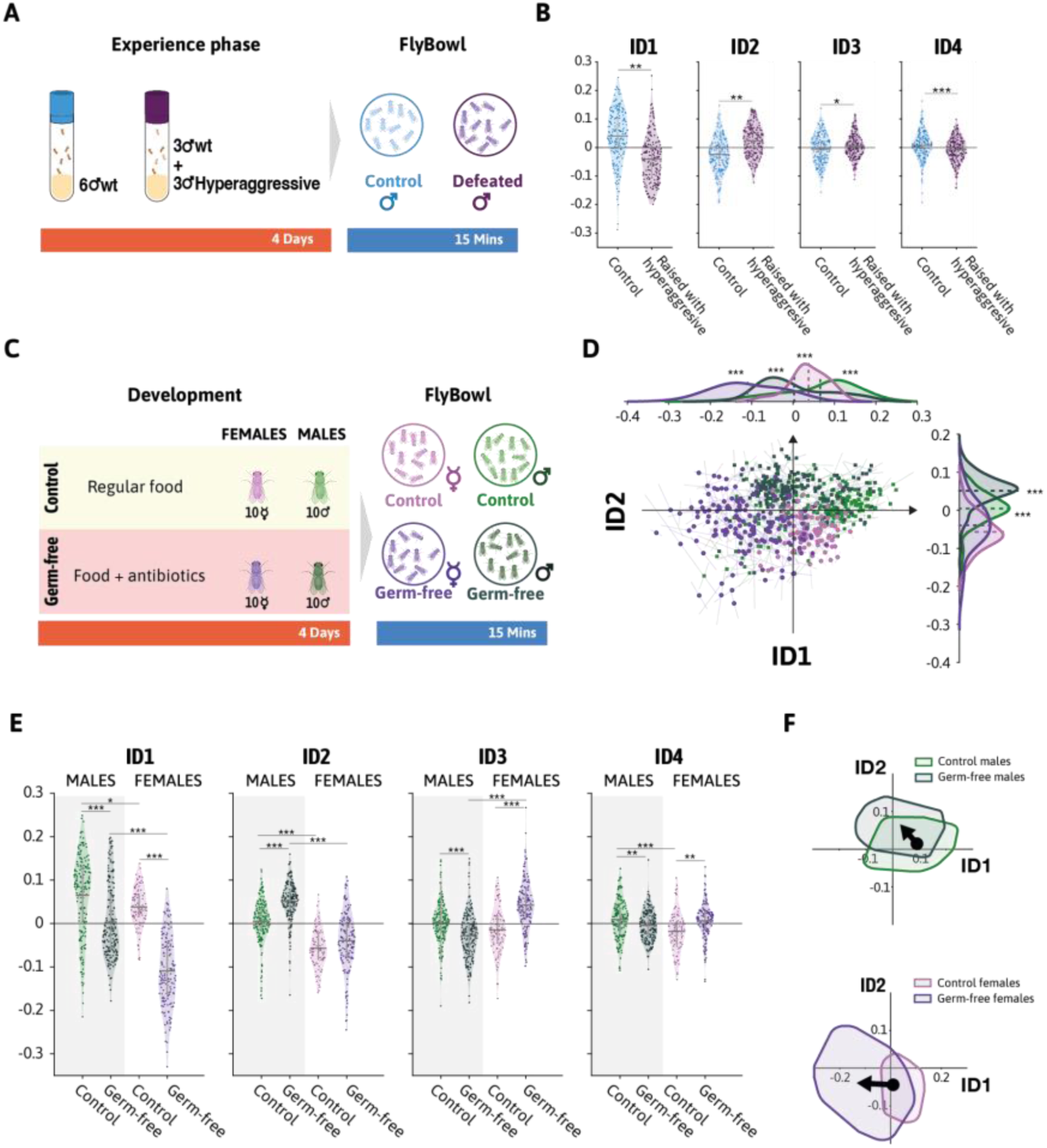
Microbiome and Social stress shape personality trait dynamics in Drosophila. Statistical significance is indicated by *, **, and ***, for p < 0.05, p < 0.01, and p < 0.001, respectively. (A) Experimental design for assessing the impact of aggressive cohabitation on identity traits. (B) trait differences of flies raised with hyper-aggressive flies (purple) compared to the control group (blue). This panel shows violin plots illustrating the distributions of ID values for control (N=200, light blue) and flies raised with hyper-aggressive flies (N = 200, purple). One one-way permutation test was performed to assess differences between conditions. (C) Experimental design for exploring microbiome influences on behavioral identity traits. (D) behavioral space of the prominent IDs. (E) trait differences of flies from both sexes raised on food with antibiotics (axenic) (females: dark purple N=151, males: dark green N=190) compared to the control group (females: light purple N=190, males: light green N=90). This panel shows violin plots illustrating the distributions of ID values. A one-way permutation test was performed to assess differences between conditions. (F) Force map of trait shifts. This panel depicts the directional shifts of clusters from the control group (males or females) to the axenic condition using arrows. The arrows represent the “forces” driving trait changes due to physiological experiences.

After exploring the influence of early-life social experiences on behavior, we next examined how physiological factors shape trait expression. To this end, we manipulated the gut microbiome, a known modulator of both physiological and behavioral responses across species, including *Drosophila* (Cryan and O’Mahony, 2011; Neuman et al., 2015; DeYoung et al., 2016; Silva et al., 2021; Schretter et al., 2018). Alterations in microbiome composition have also been linked to mood disorders and anxiety, highlighting the gut-brain axis as a critical pathway influencing personality (Clapp et al., 2017). To investigate whether the microbiome affects an individual’s position within the behavioral trait space, we reared flies from embryogenesis to adulthood on food supplemented with a cocktail of antibiotics, producing microbiome-depleted (germ-free) flies (Fig. 4C, Supp. Fig. 3C). We then compared the distribution patterns of socially raised male and female germ-free flies to their control counterparts. In the first trait dimensions, germ-free males and females both showed reduced activity scores (ID1) and increased sociability scores (ID2), suggesting a consistent microbiome-dependent effect across sexes (Fig. 4D-E). The reduced scores in space-use were not due to inherent differences in activity levels or dysregulated locomotor responses resulting from antibiotic treatment, as axenic flies exhibited normal climbing activity (Supp. Fig. 3C). This suggests that the observed differences emerge specifically from interactions within groups. Interestingly, significant sex-specific differences emerged in ID3 and ID4: germ-free males exhibited lower scores in both axes, whereas germ-free females showed elevated scores in these same traits. This divergence is particularly notable, as it suggests that germ-free females occupy higher positions on both traits that are often behaviorally linked but regulated differently in the absence of a microbiome.

Lastly, force map analysis of intersectional trait spaces further supported these observations (Fig 4F). In both sexes, germ-free flies shifted toward lower activity (ID1) and higher social investment (ID2). However, dimorphic patterns became evident when ID3 and ID4 dimensions were included: males and females exhibited opposite shifts, with the most pronounced changes observed in females (Fig. 4E). Together, these findings illustrate how social, sexual and physiological conditions dynamically modulate individuality and behavioral traits, with sex-specific effects and the interplay of life experiences and physiological changes acting as key drivers of variation. Our findings demonstrate that the trait space defined by the four IDs effectively differentiates individual flies, assigning each a unique combined score. Furthermore, social, sexual and microbiome influence the positioning of individuals within the trait space while preserving differences among individuals within the same cohort.

## Discussion

In human psychology, we often distinguish between *temperament*, the innate, biologically rooted tendencies to respond to the environment, and *personality*, which builds upon temperament through life experiences and learned values (3). Here, we investigated how life experiences shape personality-like traits in *Drosophila melanogaster*.

Using the previously established Identity Domains framework, a unified analytic approach for studying personality across species, we identified four trait dimensions: space use, time use, and two additional axes. The latter two are provisionally associated with avoidance and aggression, but their exact interpretations require further investigation. Notably, we uncovered pronounced sex-specific differences in how these traits shift in response to various life events, including social isolation, mating experiences, stressful encounters, and microbiome manipulation. Thus, showing each fly’s behavior is the product of a conserved foundation modulated by its unique life history.

These results parallel the phenomenon of polyethism in social insects, where age and experience shift the likelihood of performing particular tasks, such as nursing or foraging, along constrained behavioral axes defined by the needs of the colony (78). Analogously, in *Drosophila,* we observe systematic, experience-dependent displacements within identity space that preserve a stable baseline (temperament) while rebalancing behavioral modules (personality). Thus, as in polyethism, experience channels behavior within an evolutionarily “permitted” manifold rather than generating unconstrained novelty. The key distinction is the source of selection: in eusocial insects, trajectories are coordinated for colony needs, whereas in flies, they manifest as individual personality change with direct fitness consequences. These parallels establish identity space as a shared language for comparing colony roles in eusocial insects with individual personalities in solitary species, facilitating a mechanistic understanding of experience-driven shifts within a conserved behavioral geometry.

Another key finding from the identity space framework is that flies respond differently to familiar and unfamiliar social environments. Flies reared in groups scored higher on ID2, the axis representing time invested in social interaction, when paired with unfamiliar partners. This pattern resembles the “social novelty” effects observed in mammals, where interactions with new individuals lead to heightened investigatory behavior (79). Similar forms of discrimination are also found in eusocial insects, where social behavior is organized around colony-level recognition systems. In these systems, individuals rely on chemical cues to distinguish nestmates from outsiders, enabling cooperation within the colony and defense against intruders (80, 81). *Drosophila* lacks such a colony-based structure, making the observed differences particularly intriguing. These differences could reflect recognition at the level of individual flies, but they might also arise from changes in group composition or shared social environment. In this view, novelty in the social setting, such as a shift in group membership or even social “culture” (82), can increase social interaction without requiring explicit identification of particular individuals.

Our study enriches the ongoing discourse about genetic and environmental contributions to behavioral individuality. Drawing parallels to Freund et al.’s work with genetically identical mice (9), our findings underscore the profound role of non-genetic factors in shaping individual differences. Like the mouse model, *Drosophila* exhibited consistent individual differences despite genetic similarity and controlled environmental settings.

Another novel aspect of our study was the integration of large language models (LLMs) to objectively assist in interpreting behavioral traits. Traditional trait labeling in behavioral science can suffer from subjective bias, often relying heavily on researchers’ intuition. By employing multiple state-of-the-art LLMs, including OpenAI’s ChatGPT, Google’s Gemini, and Anthropic’s Claude, to independently interpret correlations between our measured behaviors and derived traits, we significantly reduced human bias and enhanced interpretive transparency. The models showed strong consensus on certain trait labels (e.g., “activity” and “sociability”), while identifying more ambiguous behavioral categories in others. This innovative use of LLMs not only validates our methodological approach but also illustrates the potential for AI-driven tools to bring greater objectivity and reproducibility to personality research across species.

Behavioral individuality in *Drosophila melanogaster* reflects a fundamental evolutionary balance between selection for optimal trait values and the need to maintain variability for unpredictable environments. This concept of bet-hedging suggests that populations hedge their bets by preserving diverse phenotypes to enhance survival in changing conditions (6, 83, 84). In social contexts, individual differences stemming from unique interaction histories contribute to inter-individual differences in group dynamics and behavior. In our study, we found distinct coping mechanisms that accompany life events, often in a sex-specific manner. We further found that early-life experiences did not produce unconstrained behavioral novelty, but instead shifted individuals within a limited, structured trait space - suggesting that phenotypic plasticity generates adaptive diversity along evolutionarily “permitted” axes of variation.

Previous studies using isogenic flies raised in uniform environments have shown that individuality is far from random and animals consistently vary along a small set of covarying behaviors, forming robust behavioral syndromes (62, 63, 85). High-throughput tracking revealed sparse but significant trait covariances that populate a multidimensional personality space, with the magnitude of intragenotypic variability modulated by genotype, behavioral domain, and environmental complexity (66, 86–88). Individuality even extends to associative learning, where flies retain stable idiosyncrasies across conditioning paradigms. The present approach builds on this foundation in two ways. First, by continuously tracking animals while they interact in group settings, we quantify individuality in a genuinely social context rather than in isolation. Second, we analyze these data with the Identity-Domain (ID) framework, which advances the behavioral-syndrome concept by explicitly linking between-individual variability to within-individual stability and by distilling behavior into a concise, interpretable set of trait dimensions. Together, these advances reveal how social experience, mating history, and microbiome state reposition flies within a stable, low-dimensional space of personality-like traits.

We found that activity levels and sociability were significantly modulated by early life experiences, with distinct responses observed in male and female flies. Social isolation generally increased activity levels as well as ID4 (aggression) scores, suggesting heightened exploratory and competitive behaviors, whereas mating reduced activity and exploration but increased time investment in social interactions, particularly pronounced in males. Interestingly, ID3 (avoidance scores) exhibited complex sex-dependent patterns, with mating markedly increased avoidance in males, whereas social isolation primarily affected females. The observed sex-specific responses underscore a fundamental sexual dimorphism in how experiences shape behavior, particularly along the social interaction axis. Socially isolated males became more socially interactive, whereas isolated females showed reduced sociability, suggesting distinct adaptive strategies between sexes. The clear impact of social isolation on female personality-like traits underscores the analytical power of the identity domains, which can resolve differences between conditions previously indistinguishable with traditional behavioral metrics.

Other conditions also significantly affected trait expression. Prior experience with aggressive conspecifics led to increased social time investment and decreased aggressive territoriality in socially defeated male flies. This contrasts with rodent studies, where social defeat typically induces social avoidance (89), suggesting that male flies may adopt sociability as a compensatory strategy in stressful social environments. The mechanisms behind this unexpected increase in sociability remain unclear, suggesting complex, context-dependent behavioral adaptations to social stress. The microbiome also emerged as a crucial modulator of behavioral traits, influencing the expression of the traits associated with aggression and avoidance differently across sexes. Germ-free flies displayed reduced exploratory behavior yet increased social interactions, with notably dimorphic responses.

Germ-free males exhibited lower scores in both ID3 (avoidance) and ID4 (aggression), while germ-free females showed elevated scores in these same traits. This divergence suggests that germ-free females occupy higher positions on both avoidance and aggression axes, traits often behaviorally linked but regulated differently in the absence of a microbiome. These microbiome-dependent shifts, observed specifically within social contexts rather than general locomotor activity, reveal the microbiome’s profound role in shaping socially relevant behaviors, possibly via alterations in pheromone signaling or sensory perception (90). These findings are not surprising considering the multifaceted roles of microbiome composition on the physiology and behavior of flies, including effects on activity (91), pheromone production (90, 92, 93), aggression (94), and mating-related behaviors (95).

Another interesting aspect of our findings is found when combining two identity domains. For instance, the coupling of sociability and aggression scores can be used to define the valence of social encounters. Since both solitary and mated male flies showed high sociability scores, but solitary males exhibited significantly higher agonistic scores, suggesting that while both conditions promote interaction motivation, the nature of interaction in solitary males is more competitive/aggressive. The co-regulation of sociability and agonistic behavior was also evident in germ-free male flies. These findings underscore how different forms of social deprivation trigger distinct behavioral responses, emphasizing the importance of context in shaping social behaviors. The coupling is weaker in female flies, where agonistic behavior scores are relatively low even when sociability is high (e.g., mated females), suggesting this dimension reflects dimorphic differences in social behavior.

While this research demonstrates the presence of personality-like traits in *Drosophila*, several limitations should be considered. First, the relatively short duration of behavioral recordings (15 minutes) restricts our ability to fully assess trait consistency over extended periods. Nevertheless, the observed stability of identity domains across varying social contexts, such as interactions with familiar versus unfamiliar individuals, supports their validity as robust, personality-like traits. Second, personality studies typically evaluate trait consistency across multiple contexts or tasks. In this study, trait assessment was based on a single behavioral paradigm; therefore, further research using additional behavioral contexts is necessary to confirm whether individual trait scores remain consistent across different scenarios. Yet, most personality studies use a limited number of behavioral measures to discriminate between individuals, and therefore although we tested the flies in one task, the data obtained from the FlyBowl setup is very rich, and together with the dynamic nature of social interaction that in which each interaction affects the outcome of future ones, provides an intricate and holistic view of individual traits.

In conclusion, our study demonstrates that behavioral individuality in *Drosophila* is both deeply rooted and dynamically malleable. Flies possess stable, consistent behavioral tendencies akin to temperament, yet those baseline traits can be systematically altered by social interactions, mating experiences, and even gut microbiota composition. This interplay of constancy and change yields a distinctive behavioral profile for each individual, effectively a personality, even in an organism with a relatively simple brain. By integrating evolutionary theory with modern high-dimensional behavioral analytics, we highlight how life history can sculpt individuality beyond an animal’s genetic blueprint. The introduction of the Identity Domain framework was key to uncovering these effects, and it provides a blueprint for quantifying personality-like traits in other models. Together, our findings offer a unified view of personality as the product of inherited dispositions and accumulated experiences, a view that transcends the human condition and extends to the smallest of animals. We propose that the biological logic of personality, revealed here in flies, follows general principles that apply across the animal kingdom. Elucidating that logic in simple model organisms paves the way toward a deeper understanding of the origins of personality in more complex species, including ourselves. Ultimately, this work moves us closer to unraveling how nature and nurture conspire to shape the spectrum of individual behaviors that we recognize as personality.

## Materials and Methods

### Fly Lines and Culture

*Drosophila melanogaster* WT Canton S flies were kept at 25C,° 60–70% humidity, light/dark of 12:12 hours, and maintained on cornmeal, yeast, molasses, and agar medium, unless otherwise specified. Behavioral experiments were conducted on 4-day-old, ∼1.5 hours after lights on. Cyp6a20-GAL4 was obtained from the Heberlein collection, and Cyp6a20-RNAi was obtained from VDRC.

### Analyzing Social Group Interaction Using the FlyBowl System

Group behavior was analyzed using the FlyBowl system. Ten flies were inserted into each FlyBowl arena, and their behavior was recorded for either 15 or 30 minutes, followed by tracking using CTRAX (Kabra et al., 2013). Tracking errors such as identity merging, misidentification, and discontinuous trajectories were fixed using FixTRAX (Bentzur et al., 2021). Following tracking, behavioral classifications were performed using JAABA (Janelia Automatic Animal Behavior Annotator; Kabra et al., 2013). Quantitative analysis and time-series visualizations were generated using MATLAB scripts incorporating JAABA Plot (Kabra et al., 2013). Experiments exhibiting unresolved tracking errors were excluded from downstream analysis to ensure data integrity.

### Behavioral Ethograms

Ethograms were designed to capture the temporal structure and sequential organization of behaviors exhibited by individual flies within each experimental group. Per-frame behavior annotations were extracted from JAABA output files and depicted as binary vectors for the presence or absence of nine distinct behaviors across all video frames. The resulting matrices were visualized as heatmaps, with time (in minutes) on the x-axis, behavior identity on the y-axis, and color-coded segments indicating behavioral expression.

### Aggregated Behavioral Timelines

Aggregated timelines were used to compare the overall behavioral profiles across experimental conditions. For each fly, the relative proportion of time spent performing each behavior was calculated based on the total time during which any behavior was detected. As with the ethograms, a binary behavior matrix was generated, with rows representing behaviors and columns representing time points. The total duration of each behavior was obtained by summing the entries in each row, yielding the number of frames during which each behavior occurred. These values were normalized by the total number of behavior-expressing frames (i.e., the sum of all row totals), then converted to percentages.

### Manipulating Social and Sexual Experiences

Wild type (wt; *Canton S*) flies were anesthetized with CO₂ shortly after eclosion and placed in food vials under one of three conditions: isolated (single sex), grouped (10 same-sex flies), or mixed (5 males and 5 females) for four days under a 12 h light:12 h dark cycle. On the fourth day, isolated flies were combined into same-sex groups of 10, and mixed-sex flies were separated by sex and combined with same-sex flies from other vials. Flies were inserted into the FlyBowl arenas in groups of 10 and allowed to habituate for 1 minute before testing. Behavior was recorded for 15 minutes.

### Testing Group Familiarity

Male flies were socially raised in groups of ten and were divided during the test into two cohorts: one tested with familiar flies (with whom they grew up) and the other tested with socially raised flies from other groups (each fly derived from a different vial). Behavioral tests were conducted under standardized conditions to minimize environmental variability (Bentzur et al., 2021).

### Generating Socially Defeated Flies

Wild type (wt) and *Cyp6a20-GAL4/+; UAS-Cyp6a20-RNAi/+* male flies were collected within two hours of eclosion using light CO₂ anesthesia and housed in groups of six per food vial. For the experimental group, three *Cyp6a20* knockdown (kd) males and three wt males were housed together in glass vials (10 × 75 mm) containing standard food and maintained at 25°C with a 12 h light:12 h dark cycle for four days. Control groups were composed of six wt males that were housed under identical conditions. At the end of the experience phase, wt males housed with *Cyp6a20* K.D males were collected and pooled into groups of ten flies. Control wt males were similarly pooled into groups of ten. All groups were transferred to FlyBowl test arenas in batches of ten flies per arena under standardized loading conditions. Flies were allowed to habituate in the arenas for one minute before behavioral testing commenced.

### Generation of Germ-Free Flies

Wild-type (WT) flies were reared in bottles containing 50 mL of standard fly food supplemented with a cocktail of antibiotics: 50 μg/mL tetracycline, 200 μg/mL rifampicin, and 100 μg/mL streptomycin. Behavioral assays were conducted using the F1 progeny. Shortly after eclosion, F1 flies were anesthetized with CO₂ and assigned to either the antibiotic-treated or control group. Control flies were maintained on untreated standard food. Both groups were sex-separated and housed in vials (10 flies per vial) under a 12 h light:12 h dark cycle for four days. On the fourth day, flies were transferred to FlyBowl test arenas in groups of 10 and allowed to habituate for 1 minute prior to behavioral testing. To evaluate the efficacy of the antibiotic treatment, DNA was extracted from three wt flies reared on standard food and seven wt flies reared on antibiotic-supplemented food. DNA extraction was performed using the DNeasy Blood & Tissue Kit (Qiagen, Cat. No. 69504), following the manufacturer’s protocol. Flies were homogenized in 200 μL of lysis buffer (20 mM Tris [pH 8.0], 2 mM EDTA, 1.2% Triton X-100) supplemented with lysozyme (20 mg/mL; MP Biomedicals, Cat. No. 210083401), followed by incubation for 90 minutes at 37°C. Subsequently, 200 μL of Buffer AL and 20 μL of proteinase K were added, and samples were incubated for 90 minutes at 56°C. DNA purification was completed via column-based extraction as per the kit instructions. To confirm the absence of microbial DNA, PCR was performed on the extracted DNA using universal bacterial 16S rRNA primers (16SA1: AGAGTTTGATCMTGGCTCAG; 16SB1: TACGGYTACCTTGTTACGACTT), as described by Ridley et al. (2012) with modifications from Simhadri et al. (2017). PCR amplification was carried out using KAPA2G Fast ReadyMix (KAPA Biosystems, Cat. No. KK5101) according to the manufacturer’s guidelines.

### Measuring Traits Using Identity Domains

Linear Discriminant Analysis (LDA) was used to quantify inter-individual variability in fly behavior. Behavioral data obtained from the FlyBowl system were normalized across batches, recorded monthly. For each fly, behavioral variability was analyzed over the 15-minute experiment in 5-minute segments (three segments total). LDA was applied to identify a projection matrix (W) that maximized between-cluster variability (Σb) while minimizing within-cluster variability (Σw), providing a measure of stable behavioral traits.

Flies were ranked based on pooled data per day and across segments. The relationship between behavior and fly identity was derived by solving the equation:

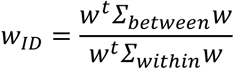

This analysis enabled the classification of behavioral traits linked to individual identity.

### Pareto Optimality

Pareto task Interference (ParTi) analysis (Shoval et al., 2012) was used to identify fly archetypes and assess personality-like behavioral dimensions. The analysis was conducted in MATLAB using the Pareto Task Inference package. Archetypal behavior was analyzed within a space spanned by identity dimensions ID1 and ID2, allowing for the visualization and interpretation of data structure.

### Fisher-Rao Separability Analysis

Fisher-Rao separability was used to compare the distinctiveness of behavioral components across flies, mice, and randomized data. Data were loaded and analyzed in MATLAB. Separability scores for flies and mice were computed directly from the loaded datasets, while scores for randomized data were averaged to provide a baseline comparison.

### Radar Plot Analysis

Radar plots were generated using MATLAB to visualize subject-specific behavioral data across three identity dimensions (ID1, ID2, ID3, ID4). Data points for each subject were calculated by projecting the W on the normalized data, with rows representing individual subjects and columns corresponding to the identity dimensions. Polar plots were created for each subject using the *polarplot* function. The angles for the axes were defined as 0°, 90°,180°, and 270°, representing the identity dimensions.

### Tree Plots

The tree plots in Figure 2 show the Pearson correlation between the IDs and the behaviors. The trees show the behaviors sorted from top to bottom by the absolute correlation coefficient. In the figures, we only show the top 15 behaviors.

### Trait Interpretation Using Large Language Models

#### Core Behaviors

To identify the behaviors that uniquely contribute to each Identity Domain (ID) while minimizing redundancy, we applied Lasso regression. This regularized linear regression approach imposes an L1 penalty on the regression coefficients, encouraging sparsity and effectively selecting a subset of informative behaviors. For each ID, we limited the number of selected behaviors to a maximum of five.

#### Using Large Language Models

To assist in interpreting the identity domains (IDs), we employed three commercially available large language models (LLMs): OpenAI’s ChatGPT 4.5, Google’s Gemini 2.5 Flash, and Anthropic’s Claude 3.7. Each model was provided with the following prompt:

> “Ignore any prior context before answering the following question.

> We measured the Pearson correlation between personality traits (labeled ID1, ID2, ID3, and ID4) and the behaviors of flies. The correlations are given in separate files corresponding to each trait. Provide a prediction regarding the label of each personality trait. Explain your reasoning.”

The models were given access to .csv files containing correlation tables between core behaviors and each ID. These files are available in the manuscript’s data repository.

### Spatial Occupancy Analysis Using Voxel-Based Heatmaps

Spatial occupancy was analyzed by dividing each FlyBowl arena into square bins and computing the 90th percentile of fly occupancy within each bin. Input data included x- and y-coordinates (in millimeters) of fly trajectories, the defined arena radius, and optional parameters for bin resolution and condition labeling. Only positions falling within the arena boundaries were considered. Fly occupancy was accumulated per frame across all individuals. For each spatial bin, the 90th percentile of occupancy values was calculated across time and flies to capture regions of frequent or high intensity use while reducing the influence of outliers. The resulting occupancy matrix was visualized as a grayscale heatmap, where higher occupancy is indicated by darker shades. Areas outside the defined arena were displayed in white.

### Self-Similarity vs. Epoch Consistency

Self-similarity and temporal consistency of behavior were assessed by comparing behavioral data across experimental conditions. Four analytical approaches were evaluated: (1) raw behavioral data (unprocessed), (2) rank-normalized data, and (3) behavioral similarity projected within fly identity space using a dimensionality reduction matrix (W). Behavioral consistency was examined over a 30-minute experiment divided into 4 equal time segments. Segments 1–2 were defined as the early period, and segments 3–4 as the late period. Data were normalized, and mean behavioral profiles were projected using the W matrix. Euclidean distances were then computed to quantify intra-fly consistency across epochs. To evaluate significance, permutation tests (10,000 iterations) were used to generate null distributions for comparison. Pairwise Euclidean distances were also calculated to determine similarity scores both within and between experimental conditions. Similarity distributions were visualized as histograms overlaid with probability density curves. Vertical markers denoted the mean within-fly distance for epoch consistency. Custom colormaps were applied to differentiate conditions, and all visualizations were formatted for clarity and uniformity.

### Statistical Analysis

The Shapiro–Wilk test was used to assess the normality of data distributions across all experiments. For comparisons between two conditions, statistical significance was evaluated using either a two-sample *t*-test (for normally distributed data) or a Wilcoxon rank-sum test (for non-normally distributed data). For experiments involving three or more conditions, one-way ANOVA followed by Tukey’s honestly significant difference (HSD) test was applied to normally distributed datasets. In cases where normality was not met, the Kruskal–Wallis test was used, followed by pairwise Wilcoxon rank-sum tests with appropriate correction for multiple comparisons. Behavioral identity (ID) traits were analyzed using one-way ANOVA combined with 1,000 permutation tests to assess differences across conditions. For significant effects (*p* < 0.05), post hoc comparisons were performed using the Tukey–Kramer method.

## Acknowledgments

This work was supported by the Israel Science Foundation (grant numbers 2505/20 and 219/24).

## Supplementary Information

**Supp. Table 1:** Definitions of behavioral features used in the LDA pipeline.

| Feature | Description |
| --- | --- |
| Grooming | Fly grooms |
| Jump | Fly jumps |
| Stop | Fly is still |
| Turn | Fly changes it moving direction |
| Walk | Fly moves |
| long approach | Fly approaches another fly from a distance |
| Short approach | Fly approaches another fly from a short distance (up to 1 body length) |
| Social clustering | Fly sits in a group with at least 3 other flies |
| prolonged contact | Social interaction that lasts more than a second |
| Touch | Fly actively touches another fly |
| Angular speed | Angular speed (rad/s) |
| Angular velocity | Angular velocity (rad/s) |
| velocity | Fly speed (mm/s) |
| Change in velocity orientation | Change in velocity direction (rad) |
| Number of close flies | Number of flies within 2 body lengths |
| Occluded field of view (anglesub) | Max FOV angle blocked by another fly (rad) in short anglesub |
| Min dist. to ellipsoid (dnose2ell) | Min distance from nose to closest fly's ellipse |
| Change in occluded view | Change in blocked FOV angle (rad/s) |
| Min dist. to center between flies | Minimum distance from focal fly center to another fly center |
| Min dist. to nose | Min distance to closest fly's nose (elipsoid to nose). |
| Min dist. to tail | Min distance from nose to closest fly's tail |
| Min dist. to center between flies change | Change in min center distance (mm/s) |
| Relative radial velocity (angle sub) | Velocity direction diff. between two flies (based on anglesub) |
| Relative radial velocity (nose2ell) | Velocity direction diff. between two flies (based on dnose2ell) |
| Diff. in orientation between two flies (anglesub) | Orientation diff. to closest (based on anglesub, rad) |
| Diff. in orientation between two flies (nose2ell) | Orientation diff. to closest (based on nose2ell, rad) |
| Angle to the closest animal | Angle to closest fly centroid (based on anglesub) |
| Angle to closest animal (dnose2ell) | Angle to closest fly centroid (based on dnose2ell) |
| Angle of focal animal (dnose2ell) | Angle from fly to neighbor ( based on nose2ell) |
| Velocity diff (anglesub) | Velocity diff. to neighbor (based on anglesub, mm/s). |
| Velocity diff (dnose2ell) | Velocity diff. to neighbor (based on dnose2ell, mm/s) |
| orientation diff to nearest fly (nose2ell) | Direction diff. to closest (based on dnose2ell, rad) |
| Velocity towards a close fly (anglesub) | Velocity toward closest neighbor (based on anglesub, mm/s) |
| Velocity towards a close fly (nose2ell) | Velocity toward closest neighbor (based on nose2ell, mm/s) |

**Supp Figure 1.**
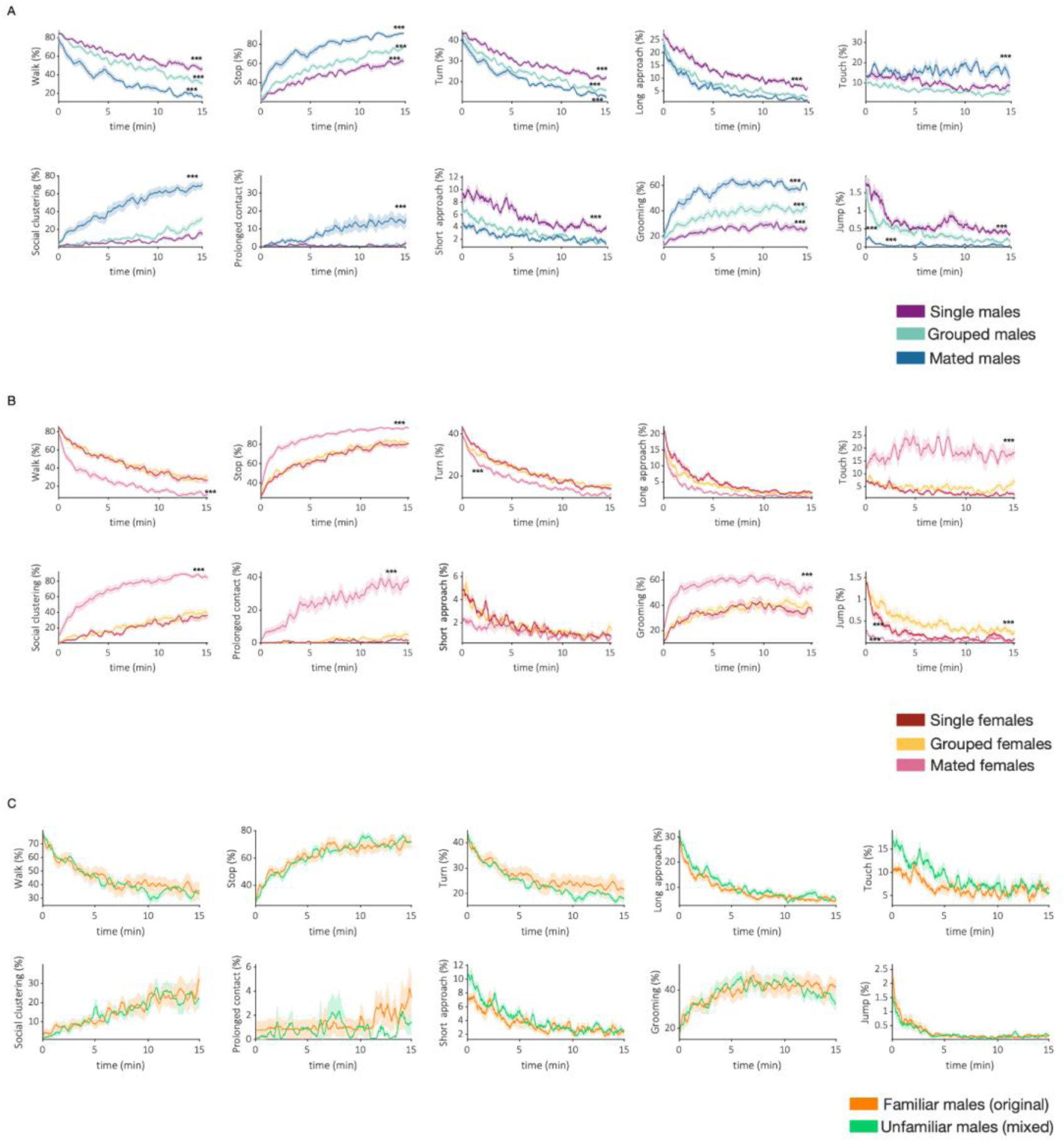
Temporal expression of selected behaviors. Average per-frame of walk, stop, turn, long-distance approach, touch, social clustering, long-lasting interaction, short distance approach, grooming, and jump behavior over 15 min of grouped, single. and mated males (A) and females (B), and familiar and unfamiliar males (C). Statistical analysis was performed on the average of each behavior for the entire duration of the test (15 min), one-way ANOVA for normally distributed features or the Wilcoxon test for non-normally distributed behaviors. FDR correction for multiple testing was performed for all analyses. Shadow signifies SEM. *** FDR<0.001, ** FDR<0.01, * FDR<0.05 for main effect.

**Supp Figure 2.**
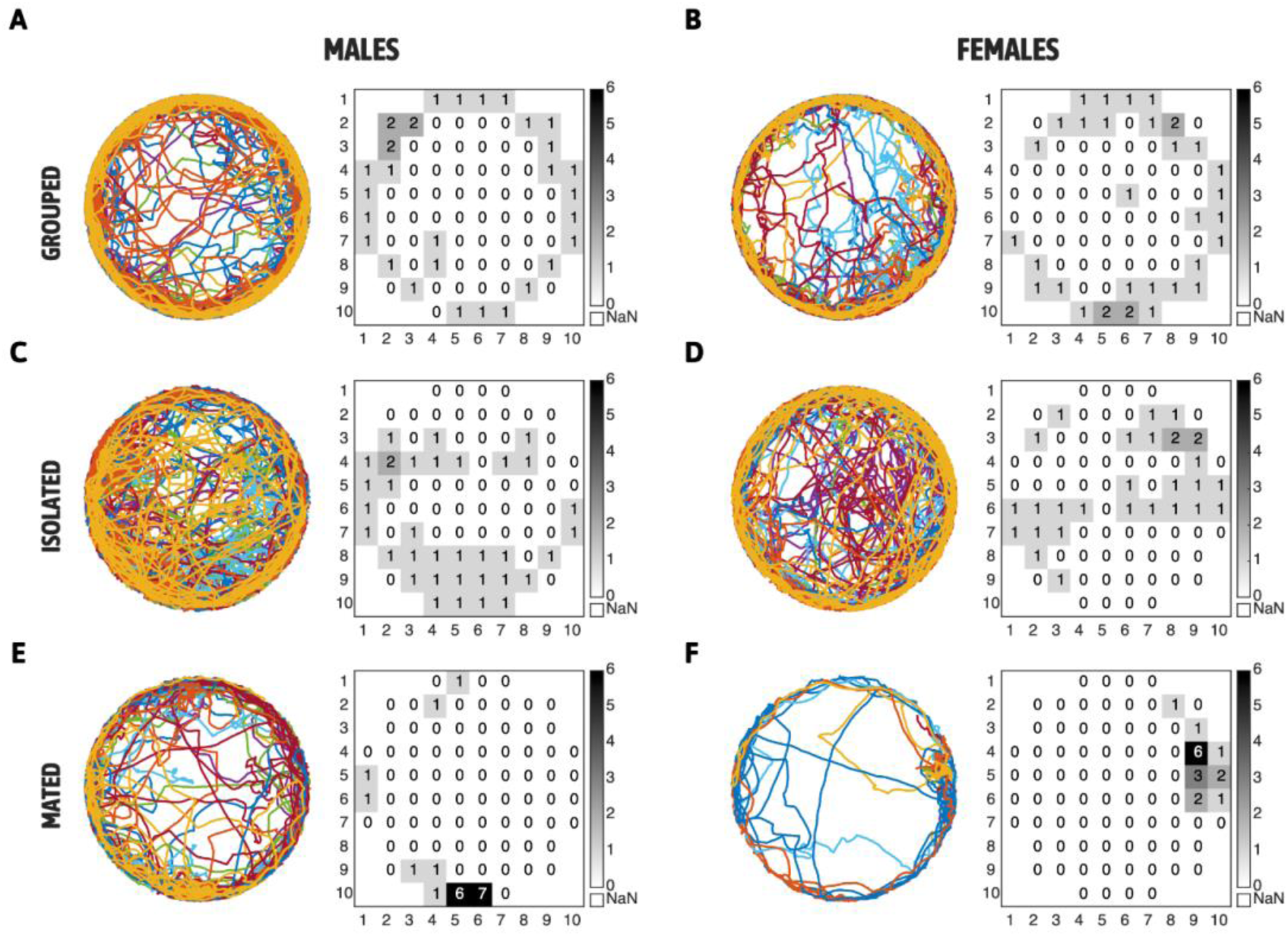
**Spatial trajectories and relative densities maps reveal sex- and condition-specific differences in exploration and social engagement**. (A–F) Trajectories (left) and spatial occupancy heatmaps (right) of representative movies in males (A, C, E) and females (B, D, F) across three conditions: grouped (A–B), isolated (C–D), and mated (E–F). Each trajectory plot shows individual paths color-coded per fly over a 15-minute session in a circular arena. Heatmaps display the 90th percentile of per-bin occupancy across flies, binned into a 10×10 grid within the arena. Gray shading indicates increasing occupancy levels (scale bar: 0–6), with white representing areas outside the arena.

**Supp Figure. 3.**
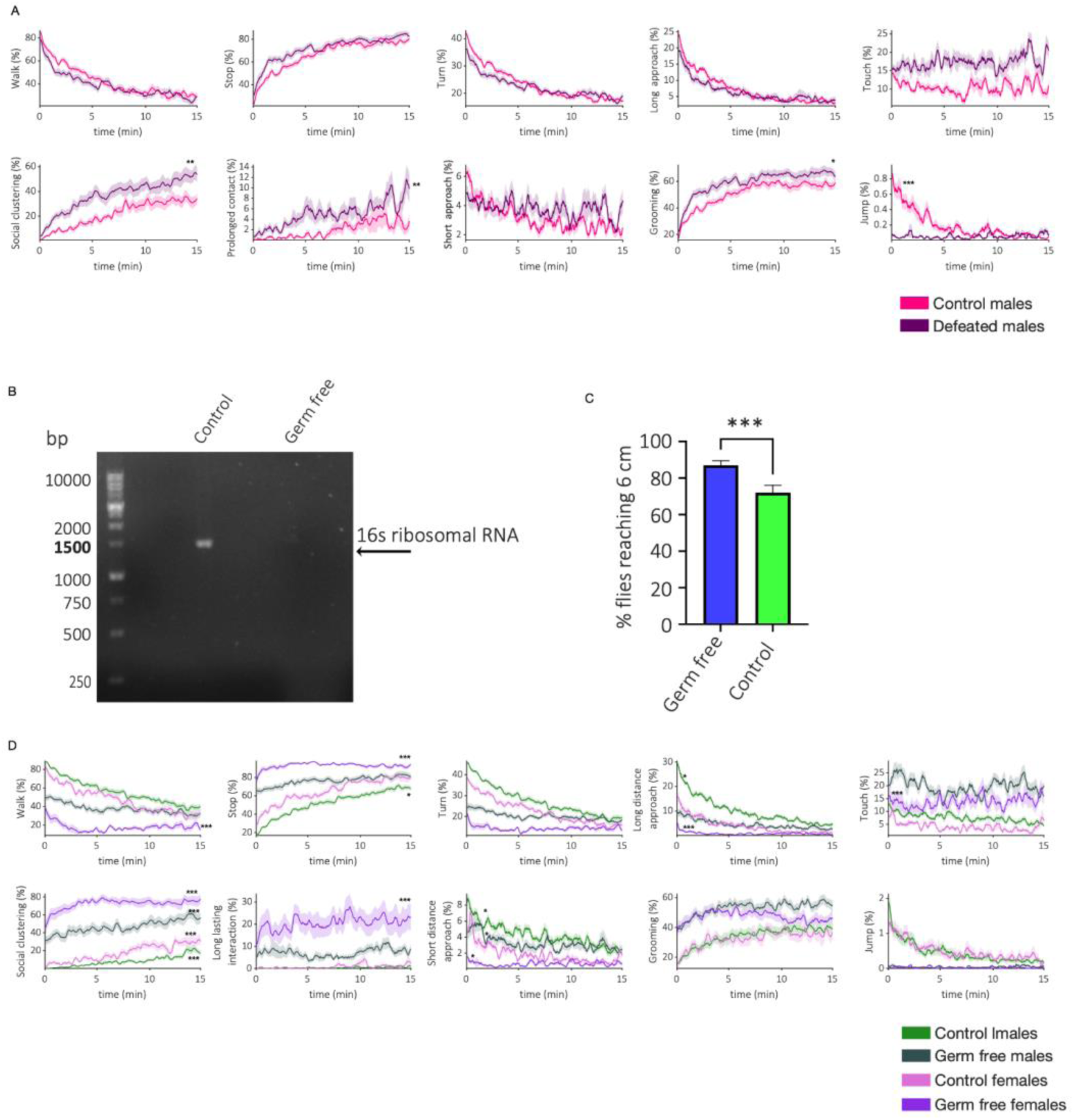
behavioral alterations and microbiome validation in control, defeated, and axenic flies. (A) Average behavioral time series of control (pink) and defeated flies (purple) (B) PCR analysis confirming microbiome depletion using bacterial 16S rRNA (∼1500 bp) as a marker, clearly observed in DNA extracted from control and not in germ-free flies. (C) Germ-free flies (blue) exhibit intact motor capacity compared to control flies (green), measured by standard climbing assay for percentage of flies reaching 6 cm threshold within 30 sec (n=10, t test ** p<0.01. Error bars signify SEM. (D) Average behavioral time series of flies from both sexes raised on food with antibiotics (germ-free) (females-dark purple N=151, males-dark green N=190) compared to the control group (females - light purple N=190, and males - light green N=90).

